## Supplementary figures and images for "Ketamine’s rapid antidepressant effects are mediated by Ca^2+^-permeable AMPA receptors in the hippocampus"

### Supplementary Figure 1

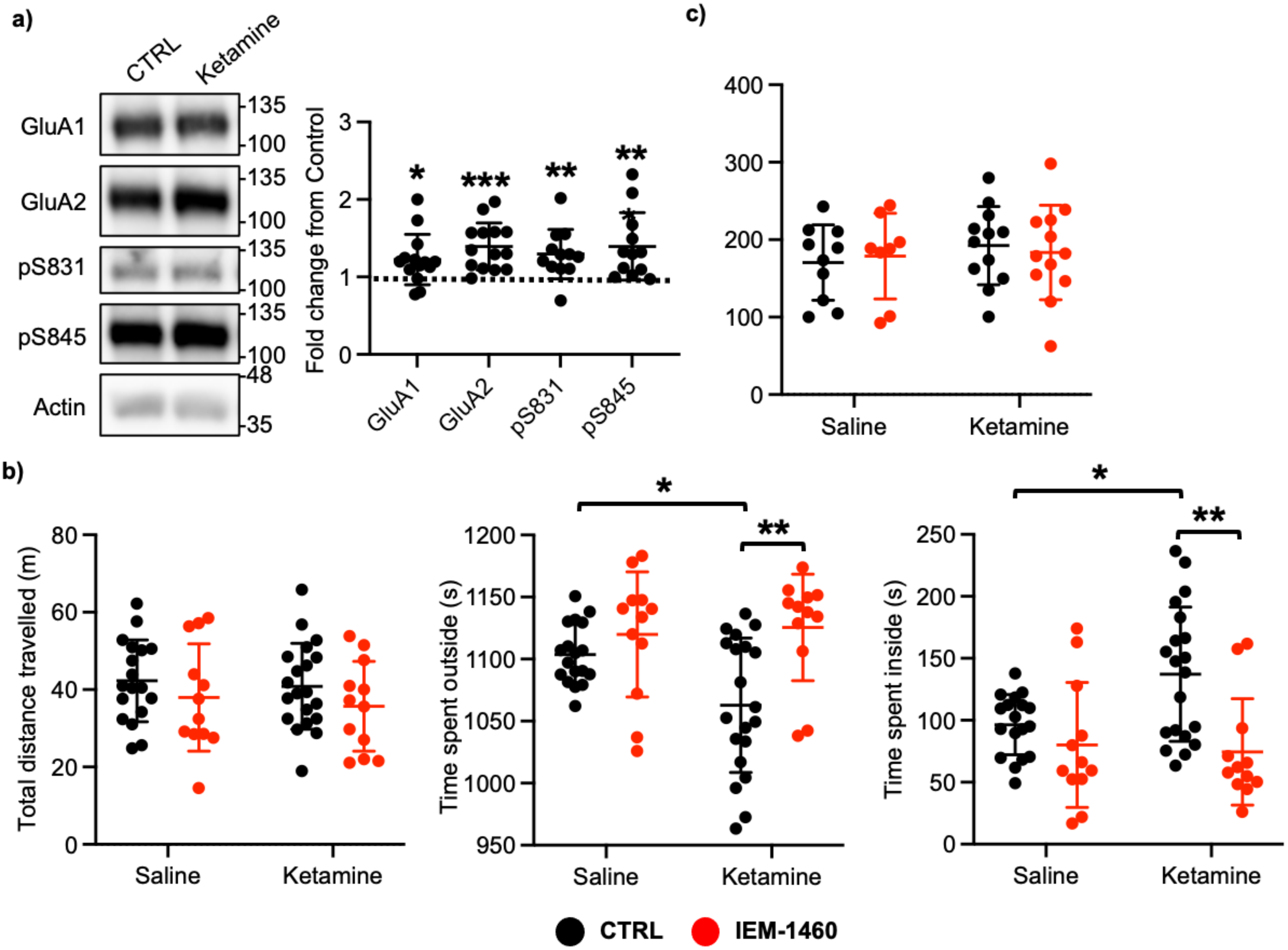
